# Faecal transplantation from exuberant toddlers increases exploratory behaviour in rats

**DOI:** 10.1101/2025.10.10.681629

**Authors:** Aatsinki Anna-Katariina, James M Collins, Munukka Eveliina, Nolvi Saara, Kara Nirit, Wilmes Lars, Deady Clara, Salin Ida, Eskola Eeva, Huovinen Venla, Lahtela Hetti, Bridgett David, Korja Riikka, Keskitalo Anniina, Isokääntä Heidi, Alex M Dickens, Lamichhane Santosh, Kallonen Teemu, Bayal Nitin, Mykkänen Juha, Pärnänen Katariina, Perasto Laura, Lahti Leo, Clarke Gerard, John F Cryan, Ted G Dinan, Karlsson Hasse, Siobhain M O’Mahony, Karlsson Linnea

**Author notes:** Corresponding author: Anna-Katariina Aatsinki. joint authorship (Siobhain M O’Mahony, Karlsson Linnea).

## Abstract

**Background:** Behavioural phenotypes have previously been transferred via faecal microbiota transplantation (FMT) from patients with psychiatric disorders to rodents. Studies indicate that the gut microbiota composition may be linked to certain temperament traits, defined as biologically-based differences in emotional reactivity and self-regulation. Here, we aimed to determine if the gut microbiota plays a role in temperament using an FMT approach. We focused on the temperament traits of exuberance, defined as positive reactivity, decreased behavioural inhibition, and high behavioural approach tendencies.

**Methods:** Faeces from 2.5-year-old toddlers from FinnBrain Birth Cohort Study with high exuberance/approach or high behavioural inhibition in the LabTAB bubbles-episode was transferred to juvenile male Sprague Dawley rats (age 22/23 days). Behaviour of the rat recipients (n=53) was assessed using the novel non-social arena, novel social arena, hole board test for exploratory behaviour, social approach-avoidance test, and forced swim test. The faecal pellets collected from the rodents were analyzed with 16s rRNA sequencing and faecal samples from the sample of toddlers (which included the donors, n=176) were analysed using short-read metagenomic sequencing. The striatum and prefrontal cortex from the rodents’ brains were analysed post-mortem using RNAseq.

**Results:** Microbiome from toddlers with high exuberance traits induced increased exploratory behaviour compared to vehicle-controls and rats receiving faeces from inhibited toddlers. Locomotor activity, social, and depressive-like behaviour remained unaffected. We noted a downregulation of the dopamine synapse pathway within the striatum of the rats that received faeces from the inhibited trait donors compared with vehicle-controls. Faecal microbiota of rats receiving faeces from the same donor resembled more each other than rats from a different cage. Clostridium species AM29 11AC in toddler microbiome was positively related to exuberance, but there were no cross-sectional associations between faecal metabolites in the human sample.

**Conclusions:** FMT from exuberant toddlers lead to altered exploratory-related behaviour in rats.

## 1 Introduction

Temperament refers to the biologically-based differences in emotional reactivity and self-regulation that can be detected during the first year of life^1^. There are arising reports on a link between the gut microbiota and temperament in infancy and childhood. Collectively, these studies have shown that gut microbiota composition is linked with an early emerging behavioural phenotype with potential relevance for the development of later psychiatric symptoms and disorders as reviewed by Alving-Jessep *et al*.^2^.

The gut microbiota is an organ-like system that is postulated to drive differences in behaviour via stimulation of the vagus nerve, production of microbial metabolites with the potential to stimulate the nervous and immune systems among other mechanisms^3^. Animal studies have shown that altering gut microbiota composition by antibiotic or probiotic treatment may result in differences in behavioural profiles^4,5^. The current understanding of the mechanisms of the gut-brain axis is mostly based on animal studies, yet there are many studies attempting to bridge the gap in human research^2^. For instance, faecal microbiota transplantation (FMT) from patients with social anxiety disorder, depression, and schizophrenia has been able to alter the behavioural patterns of the recipient animal to resemble the human condition ^6–8^. However, psychopathology often has origins in early life, and much of the previous animal work has focused on adulthood which does not consider the developmental trajectories underlying and leading to psychopathology^9^.

Temperament traits negative reactivity and fear reactivity are related to gut microbiota ^10^. Negative reactivity can be defined as behavioural withdrawal tendencies and display of negative emotions such as anger, sadness, and fear^1^. Negative reactivity and specifically fear reactivity has been a focus of temperament research given that heightened fear reactivity appears to act as a transdiagnostic phenotype predicting later internalising problems such as anxiety together with other risk factors^1,11^. From a developmental perspective, fear reactivity emerges during the second half of the first year, peaking at around 1 year of age^11^.

On the other hand, some individuals are characterised by high exuberance, which can be defined as positive reactivity, decreased inhibition, social openness, and behavioural approach tendencies^12^. High exuberance is likely based on approach-related motivation and sensitivity to reward that have neurobiological relevance^13^. As is the case with negative emotionality, high exuberance may act as a risk factor for later psychopathology, especially problematic social behaviour and impulse control problems if combined with lower emotion regulation and executive function skills^13–15^. Overall, temperament traits, in combination with other individual and environmental characteristics, relate to neurobiological and physiological differences and may increase the risk of psychopathology^16^.

Here, we aimed to investigate whether microbiota from toddlers expressing temperament extremes (exuberant or inhibited) induced changes in exploratory and activity-related behaviours in juvenile male rats. Secondly, we explored the differences in transcription profiles of the striatum and prefrontal cortex of the recipient rodents. Lastly, we aimed to screen in a well-characterised cohort of toddlers whether laboratory-tested exuberance and approach temperament traits associate with faecal microbiome composition and diversity or faecal metabolites. We hypothesised that the FMT would alter behavioural profiles of the recipient rats. Furthermore, based on earlier findings, we expected to observe associations with overall differences in gut microbiota composition, that may relate to *Ruminococcaceae, Bacteroides* and *Bifidobacterium* abundances^2^.

## 2 Methods and Materials

### 2.1 Human sample and temperament assessment

The human data was collected from the FinnBrain Birth Cohort Study based in southwestern Finland and Åland Island^17^. The study was approved by the Ethics Committee of the Wellbeing Services of the County of Southwest Finland. Written informed consent was obtained from all parents, who also provided written informed consents on behalf of their infants. The toddlers participated in Child Development and Parental Functioning Lab assessment at the child age of 30 months (± 2 months) and returned a faecal sample.

Infant exuberance was assessed with the standardised Laboratory Temperament Assessment Battery (The LabTAB), and the “Popping Bubbles” episode^18^. The episode includes a phase of low-intensity of pleasure (phase 1) and a phase of high-intensity of pleasure (phase 2). In the first phase, the child was shown how the bubbles were blown and then allowed to try themself. The second phase included three intervals where the child was instructed to pop the bubbles first, with their elbows, second, with their feet, and third, with their hands (the details of coding and reliability in Supplementary Methods).

Background variables were collected both from parental self-report questionnaires as well as linking with register data. Mothers reported the duration of exclusive and partial breastfeeding, their education level, whereas mode of delivery, neonatal and maternal perinatal antibiotic treatments, duration of gestation, maternal pre-pregnancy body mass index (BMI) and infant birth weight were collected from the hospital records. Information on received antibiotic courses were derived from the registries of the Social Insurance Institution of Finland.

### 2.2 Faecal microbiota transfer donors and sample processing

Potential donors, i.e. toddlers with high exuberance or inhibition in the “Popping Bubbles” -episode were first detected by trained psychology undergraduate students who ran the study visits. After the visit, the video recordings of these study participants (n=27) were scored by trained psychologists to confirm the phenotype. Of these 27 potentially suitable donors, 8 children were categorised as inhibited (Overall Exuberance score −1.42 - −2.26) and 10 categorised as exuberant (Overall Exuberance score 0.31-1.21). The families were contacted to recruit for the FMT study. Only toddlers with no antibiotic exposure in previous 6 months, no oral medications, no gastroenteritis in previous 6 months, and no chronic disease, such as allergies or autoimmune disease were recruited (n=12). Of the recruited toddlers, two had dietary restrictions (gluten and dairy free (excluded), avoiding mango, peach and rye). Regarding breastfeeding history, only one reported exclusive formula feeding before introduction of solid foods, and one toddler was breastfed during the sample collection, and these study subjects were excluded.

The faecal samples for the FMT study were collected from study participants homes shortly after defaecation. The duration between LabTAB assessment and anaerobic faecal sample collection was 10-123 days (median 42, mean = 52, SD 36). Fresh faecal samples were immediately transferred in pouch containing an anaerobic sachet to an anaerobic chamber, homogenised by brief vortexing with 10mM phosphate-buffered saline containing 15 % glycerol and divided into aliquots. Samples were then stored at −80°C until shipment to University College Cork, Ireland where the FMT took place.

One of the toddlers was excluded due to technical reasons. Of the 11 subjects, 4 (2 boys) had inhibited and 7 (5 boys) had exuberant temperament. For the FMT study, we selected 4 inhibited (2 boys, 2 girls) and 4 exuberant (2 randomly selected boys, 2 girls) donors.

### 2.3 Animals husbandry

All animal experiments were in full accordance with the European Community Council Directive (86/609/EEC), with an ethical approval AE19130/I046 under the licence number AE19130/I386. Male and female rats (6-8 weeks of age) were purchased from Envigo, UK. These rats were mated, 2 females to one male per cage in the Biological Services Unit, Western Gateway Building, University College Cork. After a week the male breeders were removed. Three days before birth the pregnant females were singly housed. The mothers and litters were not touched with the exception of weekly cage cleaning and ear clipping. All rat pups were weaned from the dams on postnatal day 21 and only the resulting male offspring were used as the recipient rats. Juvenile male rats (n=53 were housed 3-6 rats per cage with regular bedding material. See details on animal husbandry in Supplemental Methods.

### 2.4 Bowel cleansing and FMT procedures

The study design is presented in the Figure 1. To prepare for FMT bowel cleansing was performed, where polyethylene glycol (PEG) was orally administered 2-4 times (200-300µl) at 20-minute intervals during one day (22/23 postnatal day). All animals, including the vehicle controls, received bowel cleansing with PEG. On the day of the FMT inoculation aliquots of faecal samples were unfrozen, filtered using 50 µm filters in an anaerobic hood to remove larger particles and transferred to the animal facility on ice. Overall, there was a single freeze-thaw cycle.

**Figure 1.**
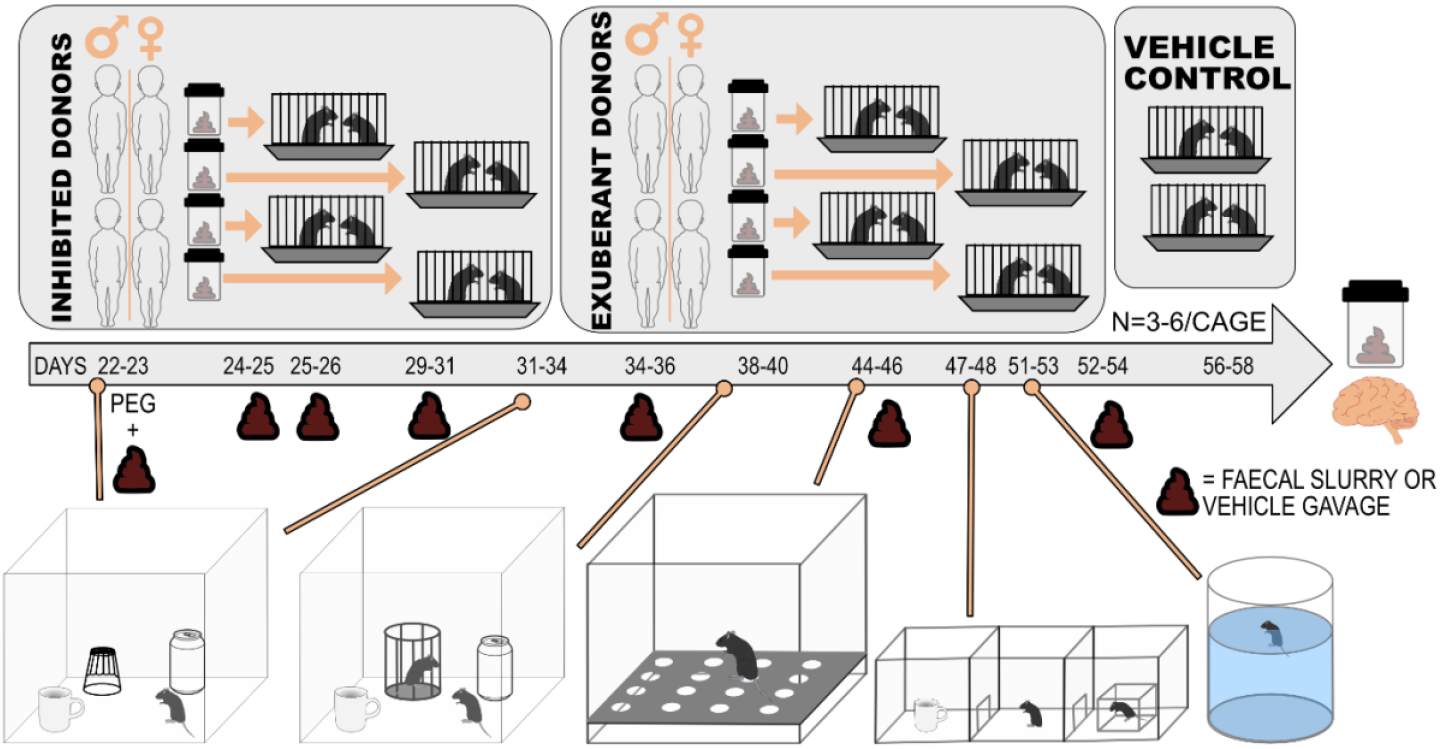
Schematic of the animal experiment timeline. The novel non-social arena (NNSA) was used at the age of 22-23 days and again at the age of 31-33 days. The novel social arena (NSA) was used at the age of 38-40. The hole board (HB) test was used at the age of 44-46 days. The social-approach avoidance test (SAAT) was used at the age of 47-49 days. The forced swim test (FST) was used at the age of 51-53 days.

Twenty minutes following the last PEG gavage, the animals received 300 μl of faecal inoculum or vehicle control (glycerol) orally (needle size). Rats in each cage received FMT from a single donor (n=4-6/rats per donor), or vehicle (n=3+6 in a cage). Rats receiving faeces from a single donor were housed in the same cage.

Animals were monitored for signs of distress and respiratory problems after each procedure. All animals received a HydroGel© sterile water hydrating gel (ClearH2O®, Portland, ME, USA) after FMT/vehicle inoculation. Two days after the initial faecal inoculation or vehicle gavage, animals received 300 μl of faecal inoculum or vehicle control boosters once a day for two consecutive days. Thereafter, animals received boosters four times with 4-8 days in between (Figure 1). There was variation in the interval between boosters to accommodate for the behavioural testing. Similarly with initial inoculation, aliquots of faecal samples were unfrozen, filtered using 50 µm filters in an anaerobic hood to remove larger particles and transferred to the animal facility on ice. Procedures are reported in accordance with the Guidelines for reporting on animal faecal transplantation (GRAFT) checklist ^19^.

### 2.5 Behavioural testing and scoring

The novel non-social arena (NNSA) was used to assess the locomotion and behavioural response to novel physical stimulus before and after inoculation (Figure 1). The novel social arena (NSA) was used to assess behavioural response to novel social stimulus (Figure 1). The hole board (HB) test was used to assess exploratory behaviour (Figure 1). The social-approach avoidance test (SAAT) was used to assess social avoidance (Figure 1). The forced swim test (FST) was used to assess anti-depressive-like behaviour (Figure 1). See Supplementary Methods for detailed description of the testing (a full description of all behavioural tests is available in the supplemental methods)

### 2.6 Sample collection

Rats were euthanised by decapitation at the age of 56-58 days (3-5 days after the last behavioural tests) and whole brains were extracted from the skull on a cold petri dish and were immediately placed on dry ice. Faecal pellets were collected from rectum, and placed on dry ice and stored at –80°C until analysed.

### 2.7 Rat brain dissection

Rat brains were removed from −80° and allowed to slowly thaw on ice. Once the tissue was slightly malleable, the striata and prefrontal cortices were rapidly dissected on ice, separated into right and left hemispheres. Brain regions were immediately placed into 200µl of RNA later and stored at −80° until shipping for RNAseq.

### 2.8 RNA extraction and sequencing of rat brain areas

The RNA extraction and RNASeq was performed by GENEWIZ (Azenta Life Sciences, Leipzig, Germany). Total RNA was extracted from tissue samples using Qiagen RNeasy Mini kit following manufacturer’s instructions (Qiagen, Hilden, Germany). RNA sequencing libraries were prepared using the NEBNext Ultra II RNA Library Prep Kit for Illumina following manufacturer’s instructions (NEB, Ipswich, MA, USA). The samples were sequenced using a 2×150 Pair-End (PE) configuration v1.5 with the Illumina NovaSeq 6000. See Supplementary Methods for detailed description.

### 2.9 DNA extraction and 16S rRNA gene sequencing from rat faecal pellets

Rat faecal pellets (approximately < 100 mg per sample, collected with 10 μl inoculation loop) were dissolved in lysis buffer and microbial DNA was extracted using a DNA Stool 200 Kit special H96 (PerkinElmer, Turku, Finland) kit with a corresponding Chemagic with Magnetic Separation Module I (MSM I) extraction robot^20^. Variable region V4 of 16S ribosomal gene was amplified with custom-design dual-indexed primers ^21^. Samples were sequenced with Illumina MiSeq (Illumina, Inc., San Diego, CA, USA) platform. Amplicon sequencing variants were inferred with DADA2 ^22^. See Supplementary Methods for detailed description.

### 2.10 Human faecal sample collection, processing and metagenomic sequencing

Regarding the faecal sample collection of the whole study sample, parents were instructed to collect the samples in a sterile collection tube as close to the study visit as possible. Moreover, parents were instructed to store the sample at +4°C and mark the date and time of sample collection. Only the samples delivered within 48 hours were sequenced (n=215 for sequencing out of 242 samples in total). DNA was extracted with GXT Stool Extraction Kit VER 2.0 (Hain Lifescience GmbH, Nehren, Germany, see details in Supplementary Methods). The starting amount of DNA was 100-500 ng. The Illumina DNA Prep Library Preparation, Tagmentation kit (Illumina) was used for the library preparation. The DNA was amplified with 5 rounds of PCR. The samples were sequenced with Illumina Novaseq 6000^21,23^ (Illumina, Inc., San Diego, CA, USA) platform for 2 x 150 base-pair reads. MetaPhlAn4 with default settings was used for profiling the microbiome composition ^24^. Overall, 197 samples were successfully sequenced and profiled. Of these, 176 had successful assessment for exuberance and approach.

### 2.11 Metabolomics of the human faecal samples

Targeted [bile acid (BAs), and short chain fatty acid (SCFAs)], and untargeted polar metabolome analysis form the human faecal samples was carried out as previously described ^25^ (Supplementary Methods).

### 2.12 Data analysis

All data analyses were conducted in R (version 4.5.1) by using packages listed below, and multiple helped functions ^26–28^. Packages vegan (2.7-1), mia (1.16.0) and miaViz (1.16.0) were used in the analyses of the microbiota data ^29–31^.

#### 2.12.1 Behavioural tests

Group differences in behavioural tests were tested with linear mixed models with cage as the random effect and the treatment group - inhibited, exuberant, and control - as the main effect with the R package nmle (3.1-164) ^32^., i.e. with formula *behavioural score ∼ behavioural group + 1*|*donor*. Since NNSA was repeated, the interaction between time and treatment group was tested to test for differences in change in locomotion, i.e. with formula *locomotion ∼ behavioural group + time of behavioural testing + time of behavioural testing * behavioural group + 1*|*donor*, where time refers to the pre- or post-inoculation timepoint.

#### 2.12.2 RNASeq

The FIMM-RNASeq data analysis pipeline was used to process the RNA sequencing data. Trim Galore was used for adapter trimming and quality filtering and STAR for aligning reads to the mRatBN7.2 reference genome from Ensembl release 111 ^33,34^. The pipeline used FastQC, MultiQC, RNASeQC, markDuplicates function from Picard followed by dupRadar, preseq, RSeQC as quality control tools ^35–40^. Subread featureCounts-function was used to count and assign reads to genes^41,42^.

DESeq2 (1.42.1) was used to identify differentially expressed genes with default options in inhibited vs controls, exuberant vs controls and exuberant vs inhibited^43^. Next, gprofiler (0.2.3) was used to test the functional pathways related to the differentially expressed genes with gSCS p-value correction and KEGG, WikiPathways and REACTOME as databases^44^. As a sensitivity analysis, a sample with > 8 % of overpresented sequences in fastQC and a sample with <60 % of uniquely mapped reads in STAR were removed from the data set, and analyses were repeated.

#### 2.12.3 Rat faecal microbiota profiles

The rat faecal pellets 16s rRNA sequencing data was agglomerated to genus level. The pair-wise distances between samples were calculated from Bray-Curtis dissimilarity. Wilcoxon rank sum test was used to test whether the distances were different if the sample was from a same or a different donor. ALDEx2 and LinDA were used to detect differentially abundant genera between treatment groups^45^.

#### 2.12.4 Human faecal microbiome and metabolite concentrations

Shannon Index, Berger-Parker index and observed richness were used as alpha diversity indices, and the correlations with temperament traits exuberance and approach was tested with Spearman correlation. Association with overall gut microbiome profile was tested with distance-based redundancy analysis using Bray-Curtis dissimilarity on relative abundance clade-level relative abundances.

Associations with species (with prevalence over 10 %) abundances were tested with MaAsLin3^46^. First, abundance associations were tested with exuberance and approach as the dependent variables without covariates. Second, interaction term between temperament and child sex was added, since gut-brain axis has been shown to have sex-specific associations. As a sensitivity analysis, subjects who had antibiotic courses during previous 6 months were removed from the analyses.

Faecal SCFA, bile acid and untargeted metabolite concentrations were log_2_-transformed with half of the minimum value as a pseudocount, and the correlations with approach and overall exuberance were tested with spearman correlation. P-values were adjusted with Benjamini & Hochberg method. The codes used in the data analysis are available in Zenodo (DOI:10.5281/zenodo.17314587.).

## 3 Results

### 3.1 Rat behaviour

#### 3.1.1 Novel non-social arena shows no difference in locomotion pre- and post-FMT

Locomotion was analysed using the novel non-social arena pre- and post-FMT. The locomotive activity, i.e. distance travelled, was not increasing differently between rats who received faeces from exuberant or inhibited toddler or vehicle-control (Fig 2A., Table S1).

**Figure 2.**
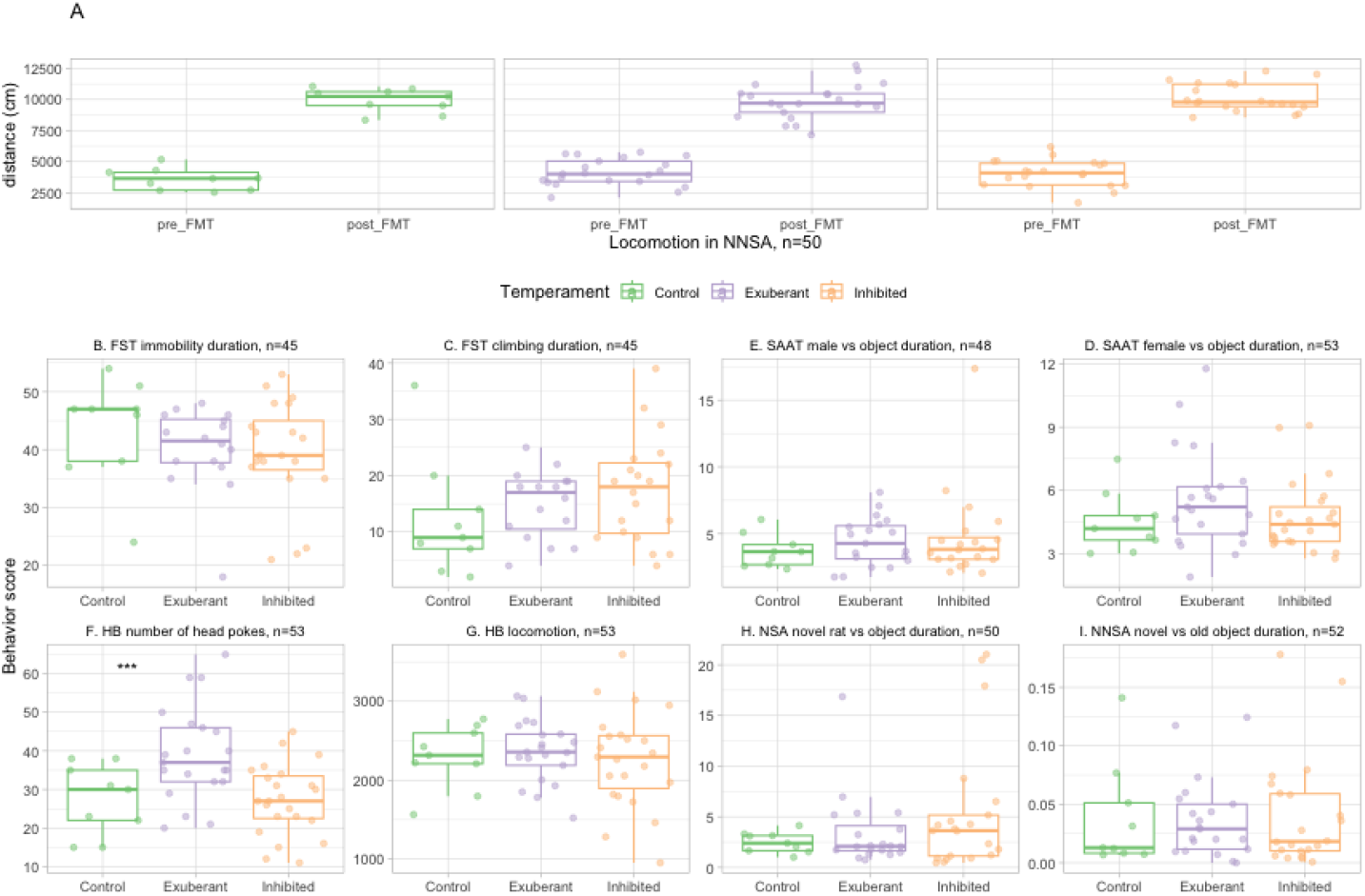
A) Change in locomotion in the novel non-social arena (NNSA) was not associated with the treatment group. There was no difference in the change of locomotion in the groups (mixed model time of behavioural assessment * group interaction term p= 0.67 for recipients of inhibited toddler’s faeces and p=0.33 for recipients of exuberant toddler’s faeces group). Panels B-I) show the treatment group differences in the behaviour, and only F) number of head pokes was higher in the recipients of exuberant toddler’s faeces group. *** ANOVA p-value < 0.005. FST = forced swim test, HB = hole board, NNSA = novel non-social arena, NSA = novel social arena, SAAT = social approach-avoidance test.

#### 3.1.2 The Holeboard test indicated higher exploratory behaviour in rats receiving faeces from exubrant toddlers but no difference locomotion

No differences were noted in locomotion during the holeboard test following FMT from the inhibited or exuberant children as compared to the vehicle control rats (Table S1). However, the rats that received the faeces from the exuberant group had an increased number of head pokes in the hole board arena compared to controls (Figure 2G., Table S1) and the recipients of faeces from inhibited toddlers (p=0.01, Figure 2G.). Controls and rats receiving faeces from inhibited toddlers did not differ (Figure 2G., Table S1). The association remained when outliers based on behavioural score were removed (> | 3 x SD|, n=34, Table S2).

#### 3.1.3 The novel social and non-social arena, social approach-avoidance test and forced-swim test were showed no difference in behavioural response to novelty or social stimulus or antidepressive-like behaviour

Anti-depressive-like behaviour indicated by crawling or immobility time in FST was not different between groups (Figure 2B-C, Table S1). Social approach or avoidance behaviour indicated by the time spent with the female or the male conspecific vs. neutral object in SAAT was not different between the groups (Figure 2E-D). Behavioural response to novel social stimulus indicated by the time spent with a novel male conspecific in the NSA was not different between groups (Figure 2H, Table S1). Time spent, i.e. duration not the distance travelled, with the novel object in the NNSA was not different between the groups that received the FMT from inhibited or exuberant toddlers or vehicle control (Figure 2I, Table S1).

### 3.2 RNA Sequencing of Striatum and Prefrontal cortex showed downregulation of genes in striata of rats receiving faeces from inhibited toddlers

The transcriptome of the striatum and prefrontal cortex were assessed in order to study potential alteration in gene expression in areas related to exploratory behaviour. DESeq2 analysis of the bulk RNASeq showed 78 downregulated and 2 upregulated genes in the striatum of rats receiving faeces from inhibited toddlers versus control rats (see volcano plot in Figure S2), whereas no genes were differentially expressed in the striatum of rats receiving faeces from exuberant toddlers nor in prefrontal cortex of any of the groups. Pathway analysis with gprofiler2 suggested that dopaminergic synpase KEGG pathway was downregulated in the rats receiving faeces from inhibited toddlers vs controls (Figure 3, gSCS adjusted p-value = 0.1, intersection size = 3 genes). In the sensitivity analysis, dopaminergic synapse pathway p-value attenuated further.

**Figure 3.**
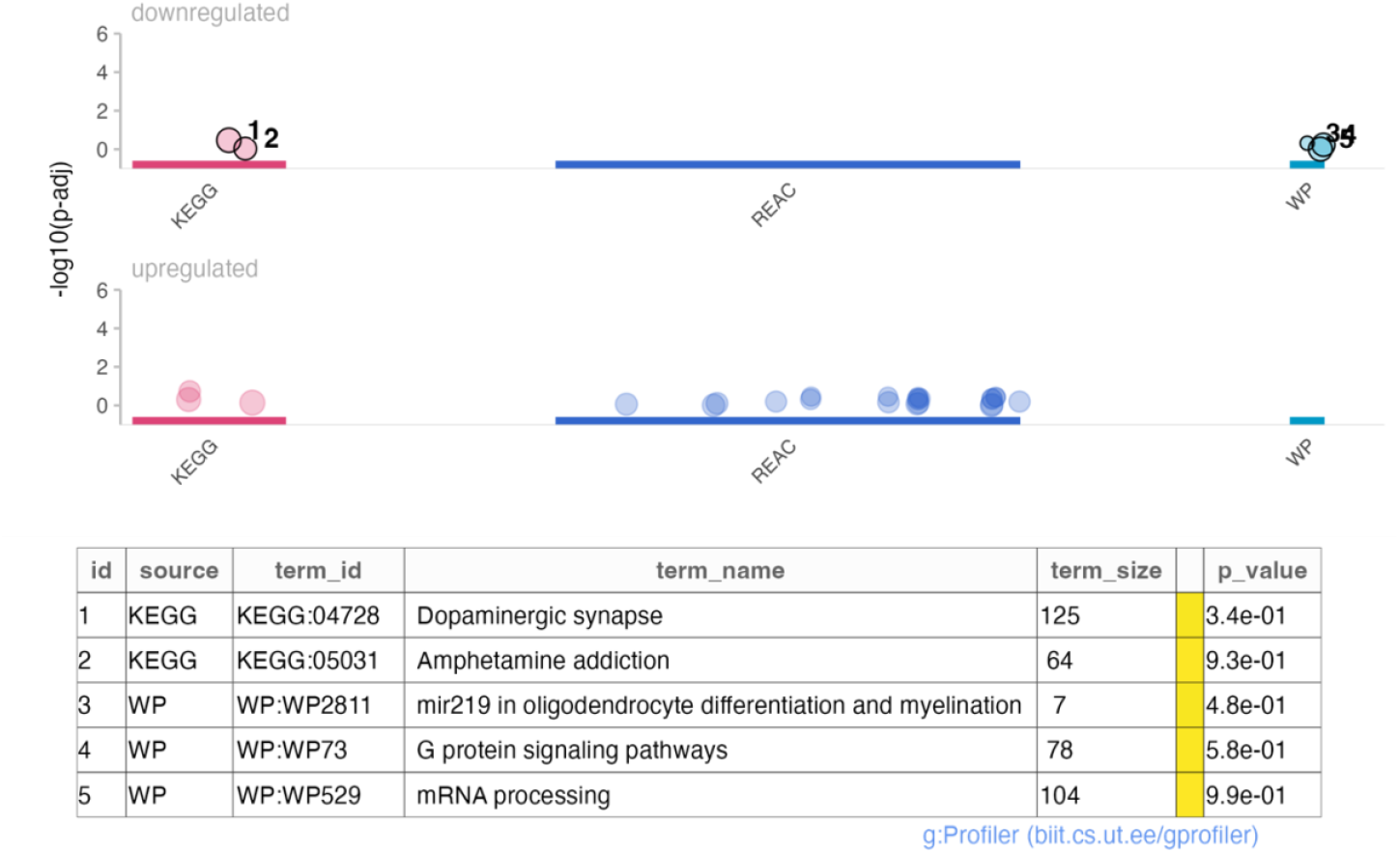
Pathway analysis of differentially expressed genes in striatum in recipients of inhibited toddler’s faeces vs controls were not statistically significant.

### 3.3 The fecal microbiota composition in rats was more cage-specific after FMT inoculation

Analysis of the microbiota revealed that the gut microbiota community composition in rat faecal pellets was more similar between rats in the same cage versus rats from different cage (Bray-Curtis dissimilarity, Wilcoxon rank sum test, p = 0.5 x 10^6^, Figure 4). However, there were no differences in community composition (distance-based redundancy analysis, PERMANOVA p=0.61) or community homogeneity (PEMRDIPS2, ANOVA p=0.67) in rats receiving faeces from inhibited or exuberant toddlers or the vehicle control (Figure 4). Moreover, there were no associations with genera abundances when correcting for multiple testing or genera presence versus absence between the treatment groups (Tables S3-S5).

**Figure 4.**
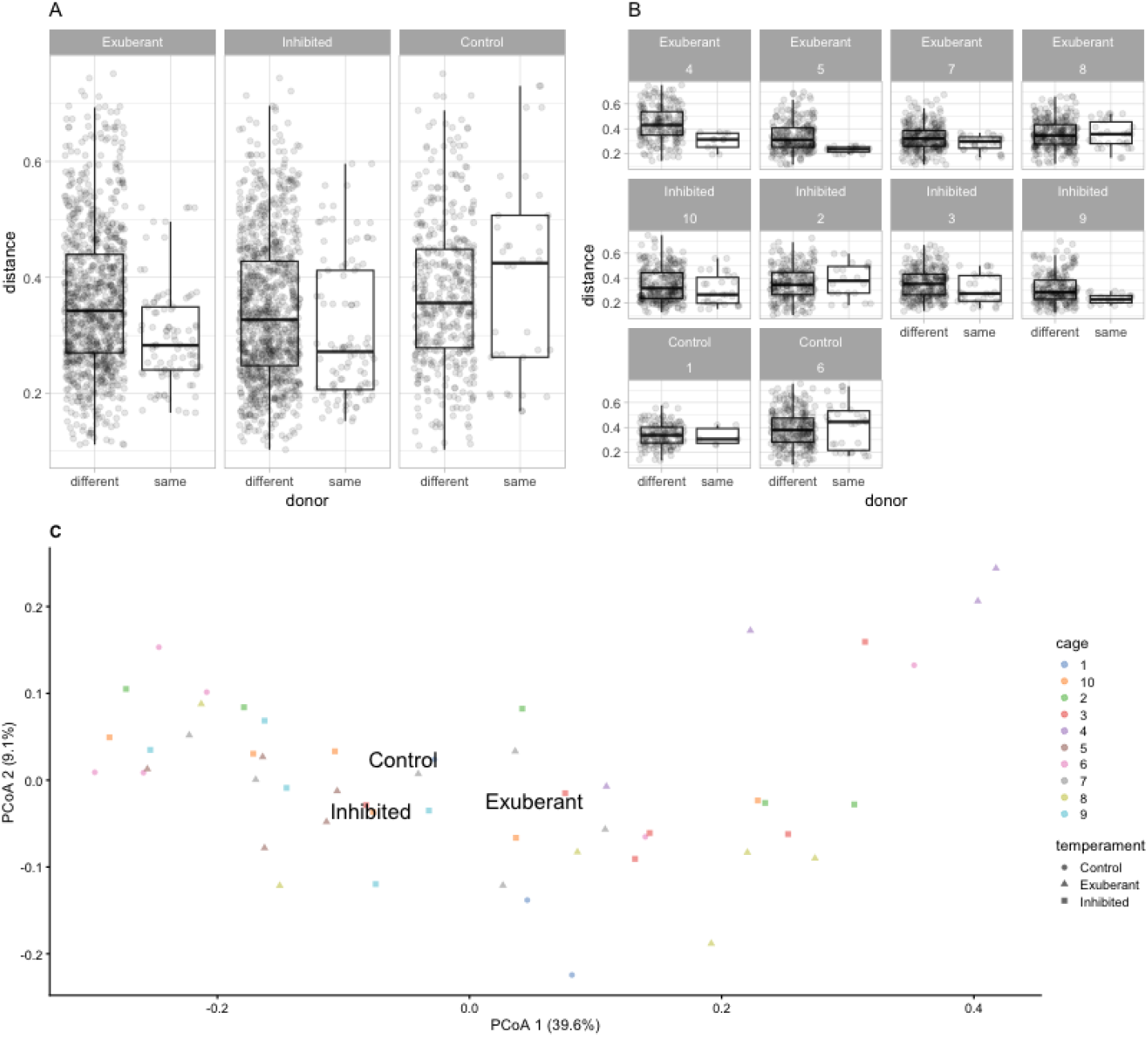
A, B) On average the gut microbiota compositon of rats from same cage resembled each other more than the rats from different cages (n=52). A) Additionally, this observation was driven by the rats receiving faecal inocula versus controls receiving vehicle. Panel B) shows similarity within a cage where the number refers to an individual cage. Additionally, the controls had higher dissimilarities to other controls than the rats in inhibited or exuberant groups (Kruskal-Wallis test p-value = 5.31 × 10^-09^, Dunn’s test adjusted p-values 2.1 × 10^-8^ for Control vs recipients of exuberant toddler’s faeces and 4.3 × 10^-8^ for Control vs recipients of inhibited toddler’s faeces). C) PCoA plot showed no clustering according to cages or treatment group. The text illustrates the center point of the group. Bray-Curtis dissimilarity was used for the dimension reduction.

### 3.4 The faecal microbiota community composition was not associated with approach and exuberance in the human sample

The whole sample of 30-month-olds with LabTAB Bubbles-episode and faecal metagenomic data available (n=172, Table S6 for sample characteristics), was used to study if gut microbiome composition is associated with exuberance and approach traits. We observed no associations between the temperament traits and alpha diversity indices (Table S7) or community composition in the whole sample or in sex-stratified analyses (Figure 5A, Table S8). In species-level in the faecal metagenomic data, none of the species were associated with exuberance when adjusting for multiple testing correction (Table S9). When removing children who had been prescribed antibiotics during previous 6 months, exuberance was associated positively with Clostridium species AM29 11AC (Table S9). Similar trend between Clostridium species AM29 11AC and approach was observed, but this was not significant after multiple-testing correction (Table S9).

**Figure 5.**
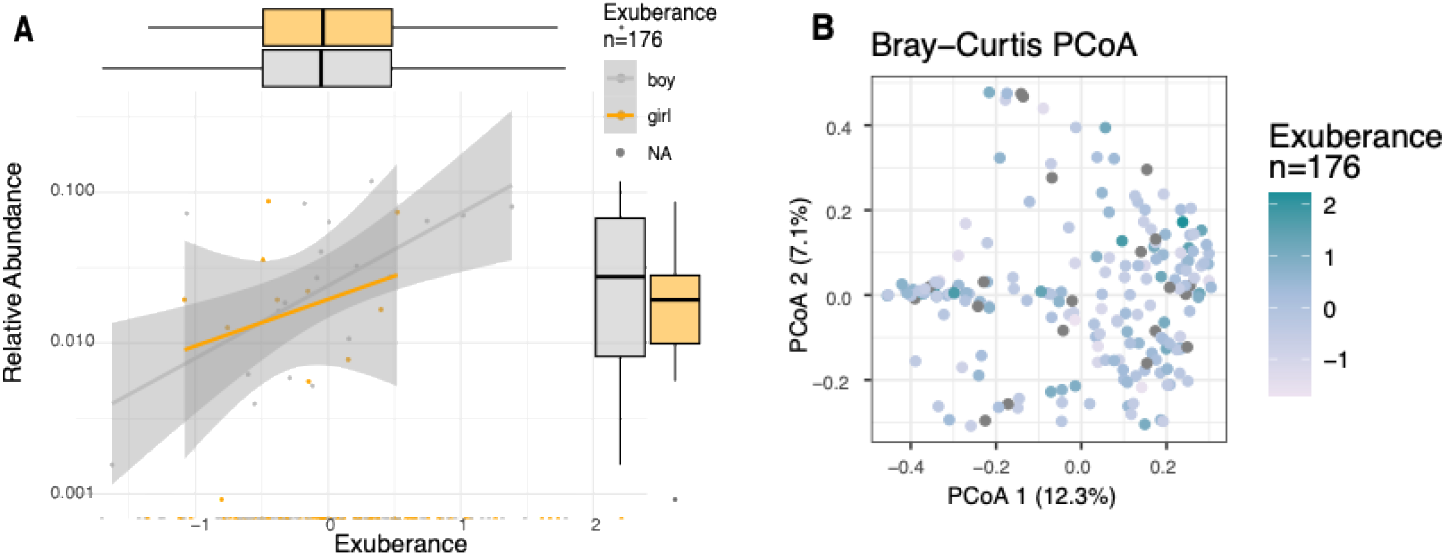
Microbiota composition and exuberance in children. A) Higher abundance of Clostridium species AM29 11AC was associated with higher exuberance. B) PCoA plot with Bray-Curtis dissimilarity on clade-level data showed no clustering according to exuberance.

Interaction between sex and exuberance or approach were associated with multiple species with nominal p-value below < 0.05 (Table S9). In sex-stratified analyses, Alistipes hominis was associated negatively with exuberance in girls, while GGB9708_SGB15236 in the family of Oscillospiraceae was associated negatively with exuberance in boys (Table S9). Abundances of Lachnospira hominis, Ruminococcus bicirculans and Waltera actigignens were positively associated with aprroch in girls, while abundances of Alistipes hominis, Clostridium species AM29 11AC, Eubacterium album were positively associated with approach in boys (Table S9).

### 3.5 Faecal metabolites were not correlated with approach and exuberance in the human sample

Temperament traits were analyzed for associations with fecal short-chain fatty acids, bile acids, and a range of untargeted metabolites, including alcohols, sugars, and energy metabolites. Our analysis revealed that UDP-glucuronic acid was positively associated with both exuberance and approach traits in the combined-sex analysis. In contrast, both exuberance and approach traits exhibited a negative association trend with sugar compounds, including D-trehalose. However, none of these associations reached statistical significance at the chosen false discovery rate (FDR) threshold of 0.05 (Table S10).

## 4 Discussion

Here we show for the first time that that that FMT from healthy toddlers presenting with exuberant temperament phenotype increases exploratory-related behaviour in juvenile rats. Furthermore, while no behavioural difference was noted in the rats receiving the FMT from inhibited toddlers, the FMT induced a downregulation of the dopamine synapse pathway in the rat’s striatum. Although we and others have previously reported associations between positive emotionality and gut microbiota composition ^2,47^, we failed to show cross-sectional associations between gut microbiome community composition or faecal metabolites and exuberance in toddlers.

Rats receiving faeces from exuberant toddlers had specifically more exploratory behaviour, but no difference in overall locomotion, depressive-like behaviour or behavioural response to physical or social novel stimulus. Exploratory behaviour is active form of novelty-seeking behaviour and relates to monoamine metabolism in the CNS, including striatum ^48^. In humans, exuberance is a phenotype that is relatively stable in childhood, and it is thought to relate to differences in the reward system, specifically approach-related motivation ^13^. Exuberance, especially when combined with other risk factors, may be linked with later impulse control problems ^13–15^. Although there were no metabolite correlations with exuberance in the human sample, rodent studies suggest that microbially produced metabolites are important for addiction, and specifically drug-seeking behaviour ^49^.

In our data, dopamine synapse pathway was downregulated in the striatum of rats receiving faeces from inhibited toddlers. Striatal dopamine activation increases exploratory behaviour and reduces self-grooming, that is a proxy for repetitive and compulsive behaviour in rodents ^50^. This aligns with the behavioural phenotype in humans, since behavioural inhibition increases the risk for anxiety disorders^51^, such as obsessive-compulsive disorder. Our observation that inhibition-related FMT could alter dopamine synapse pathway in striatum may be due to differences in intestinal metabolism ^52^. Although we could not identify candidate-metabolites in our human sample, Clostridium species AM29 11AC related to higher scores of exuberance. Earlier rodent studies have shown that gut microbiota modulation is shown to affect functioning of the reward-system and specifically dopamine metabolism ^53^. In humans, previous studies have shown that depression and quality of life relates to differences in microbial dopamine metabolite synthesizing capacity in the gut in adults ^54^. Moreover, one study showed that other monoamine pathways in faecal metagenome were positively associated with child impulsivity symptoms^55^. However, we are not aware of studies linking intestinal metabolism, potentially related to dopamine pathways, with behaviour in small children.

Although exuberant-related FMT was able to alter exploratory behaviour, we did not observe any change in the group receiving inhibited-related FMT. This may be due to behavioural test battery targeting novelty-induced, social behaviour and locomotion, while tests targeting reward-seeking behaviour, anhedonia, behavioural inhibition, repetitive/compulsive behaviour, or anxiety-related behaviour, for example might have revealed inhibition-related changes in the recipients. Alternatively, gut microbiota may influence exploratory behaviour to greater extent compared to behavioural inhibition.

### 4.1 Limitations

The current study has the merit of laboratory-based assessment of temperament. However, although laboratory-based assessment is a stable measure, the labour intensity creates limitations to the sample size of the human sample as well as the donors for FMT. This, among other factors, may have led to failing to identify bacterial profile related to exuberance. Next, available sample material in the animal study limited the duration and consequently the number of behavioural tests. Given a longer and broader follow-up, we would have been able to study, *e*.*g*. anxiety- or reward-related behaviours and other behavioural phenotypes relevant to exuberance and inhibition. Furthermore, the number of available donors was limited, which inhibited us to truly account for the modifying effect of biological sex, and the potential implication on links between temperament and microbiome ^56^.

Our study benefitted from pretreatment with PEG instead of antibiotic treatments, reducing potential confounding related to the gut microbiota eradication and unwanted central nervous system effects. While some previous studies have reported that PEG can result in alterations in the gut microbiota, bowel cleansing with PEG is an established method of achieving gut microbiota depletion of up to 90%^57,58^. Adding an additional control group with no microbiome eradication method might prove useful in future studies.

### 4.2 Conclusion

In conclusion, faecal microbiota transplantation from exuberant toddlers increased exploratory behaviour in juvenile rats. Rats receiving faeces from inhibited toddlers had downregulated dopamine synapse pathway in the striatum. Dopamine circuits in striatum coordinate reward-related behaviour, and exploratory behaviour is driven by differences in approach-related motivation^59^. Collectively, our results reveal that the reward-related features of exuberance-inhibition spectrum is mechanistically related to gut microbiota. However, we could not find a common feature relating to behaviour in gut microbiota composition or metabolome in toddlers and in the recipient rats.

## 6 Acknowledgements

Finnbrain Birth cohort Study (HK) has been funded by Research Council of Finland (grant numbers 253270, 134950), Jane and Aatos Erkko Foundation, as well as Signe and Ane Gyllenberg Foundation. Metagenomic sequencing was supported by the Jane and Aatos Erkko Foundation. AKA was supported by Yrjö Jahnsson Foundation, Psychiatry Research Foundation, Emil Aaltonen Foundation, Brain Foundation, Instrumentarium Science Foundation, Signe and Ane Gyllenberg Foundation, Duodecim Finnish Medical Society, Juho Vainio Foundation and the Research Council of Finland (grant number 347640). LK was funded by the Research Council of Finland (grant number 308176 and 325292), Yrjö Jahnsson Foundation (6847, 6976), Signe and Ane Gyllenberg Foundation, Finnish State Grants for Clinical Research (P3654), Jalmari and Rauha Ahokas Foundation, and Waterloo Foundation (2110-3601). SL was supported by the Academy of Finland postdoctoral grant (323171). LL was supported by Research Council of Finland (grant number 330887). AMD has been funded by the Waterloo foundation and Research Council of Finland (347924). “Inflammation in human early life: targeting impacts on life-course health” (INITIALISE) consortium funded by the Horizon Europe Program of the European Union under Grant Agreement 101094099 (to HK AMD).

We thank Patrick Fitgerald, Gerard Moloney, Matilda Kråkstöm, Bishwa Ghimire for their help. We thank the FinnBrain Birth Cohort Study participants and personnel, and Turku Metabolomics Center for the assistance and resources in the analyses of metabolites. Dr. Bishwa Ghimire from the InFLAMES Research Flagship Bioinformatics Team is acknowledged for creating the RNASeq data preprocessing pipeline and preprocessing the data, and for the help in interpretation of the results.

Aatsinki Anna-Katariina: conceptualization, formal analysis, investigation, data curation, writing – original draft, visualization, funding acquisition.

Collins James M, Kara Nirit, Wilmes Lars, Deady Clara, Eskola Eeva, Lahtela Hetti, Keskitalo Anniina, Pakola Ida, Isokääntä Heidi, Lamichhane Santosh, Dickens Alex, Bayal Nitin, Perasto Laura, Mykkänen Juha: investigation, writing – editing.

Munukka Eveliina: conceptualization, supervision, project coordination, investigation, resources. Huovinen Venla: writing – editing.

Bridgett David: supervision, resources, writing –editing.

Korja Riikka: supervision, resources, writing –editing, funding acquisition. Nolvi Saara: supervision, investigation, conceptualization.

Kallonen Teemu: resources, writing – editing.

Lahti Leo, Pärnänen Katariina: software, methodology, writing – editing.

Karlsson Hasse, Dinan Ted, Clarce Gerard, Cryan John: conceptualization, writing –editing, funding acquisition.

Karlsson Linnea, O’Mahony Siobhain: conceptualization, writing –editing, funding acquisition, project administration, supervision

