## Supplemental Methods and Figures for "Faecal transplantation from exuberant toddlers increases exploratory behaviour in rats"

**LabTAB Bubbles –episode**

The Lab-Tab is an assessment tool for measuring child temperament, that is designed to elicit reactive and regulatory tendencies from the children. The "Popping Bubbles" episode is designed to elicit exuberance in toddlers. The task includes a phase of low-intensity of pleasure (episode 1) and a phase of high-intensity of pleasure (episode 2). In the first episode (45 seconds), the child was shown how the bubbles are blown and then allowed to try themselves. The second episode (1 minute) included three intervals where the child was instructed to pop the bubbles first with their elbows, secondly with their feet, and thirdly with their hands. The tasks were video-recorded and scored according to the instructions of the Lab-Tab manual by trained psychologist or psychology students. Variables that were scored were as follows: a) intensity of smiling (on a scale between 0-3), b) presence of laughter (0 = no, 1 = yes), and c) vigor of approach (on a scale from 0-1 in the first episode and on a scale from 0-3 in the second episode). Reliability between two different independent coders were calculated using Cohen's Kappa for 10 of the episodes in the whole sample and were as follows; .88 for Smiling, .82 for Laughter, and .82 for Approach. For the whole episode, Cohen's K = .85.

**Animal Husbandry**

Ear clipping was done by the same researcher who conducted the behavioural testing. Dams and littermates were housed in large plastic breeding cages (45 × 28 × 20 cm), after weaning animals were housed in RC2F type cages (56 x 38 x 22 cm) in a humidity- and temperature-controlled room set to 55±10% and 21°C ± 1°C. The light/dark cycle was set to 12 hours (light phase 7am-7pm).

A Plexiglas divider was inserted between the cages holding the recipients rats receiving FMT from exuberant and inhibited toddlers to prevent any potential cross-contamination between groups. One control group was housed at each side of the Plexiglas divider. Regular rodent chow and water were available *ad libitum*. Animals were monitored daily, and body weight was measured regularly (Figure S1).

**Behavioral testing**

*Novel non-social arena*

The NNSA was a square arena covered with clean bedding material blended with bedding material from all the cages. Three neutral objects (can, mugs) were placed with even distances in the arena. The rat was placed in a ceramic bowl and transferred to the arena and was videotaped from above for five minutes. Any fecal pellets produced during the test were removed from the arena after the trial. The experiment was repeated after the inoculation and initial booster. The same arena was used with one novel object. Overall locomotion and time spent with objects (novel or familiar) were analyzed with Ethovision XT

(Noldus Information Technology, Wageningen, the Netherlands). Time spent with novel object compared to familiar object and overall locomotion in both trials were used in subsequent analyses.

*Novel social arena*

NSA was similar to NNSA, except instead of three neutral objects, the arena contained a neutral object and a novel conspecific of the same age and sex in a wire-cage. Rats were placed in the arena in a ceramic bowl and were videotaped from above for five minutes. Time spent with the object and conspecific were analyzed with Ethovision XT.

*Hole board*

The hole board was a square arena (66 x 66 cm) with an opaque floor with four holes (3.7 cm diameter, 2.2 cm deep). Three small metallic objects were placed in the holes. The animals were placed in the centre of the arena and videotaped from above for six minutes. The hole board was cleaned after each trial with 70% ethanol to remove odour cues. Locomotion in the arena was analyzed with Ethovision XT, and the number of head pokes, i.e., dipping head in the hole, was manually scored with open-source logging software BORIS (Behavioural Observation Research Interactive Software)[1].

*Social approach-avoidance test*

The SAAT arena was a rectangular arena with gray, opaque walls. The overall dimensions were 42.5 x 63 cm with two dividers with 10 cm wide openings in the middle, which resulted in three 21 x 42.5 cm compartments. The test was a three-day experiment with one habituation day and two testing days. On the first day, animals were habituated to the arena for 10 minutes. The following day, experimental animals were placed in the middle compartment, the other compartment housed a novel male conspecific of matching age in a holding unit made of clear Plexiglas with breathing holes. The final compartment had a neutral object (a mug). The rat was videotaped for 10 minutes. On the third day the experimental rats were exposed to a novel female conspecific of matching age. The time spent interacting, i.e., head towards the object or conspecific, was scored manually with BORIS. Time spent interacting with the conspecific versus the object was calculated and used in the subsequent analyses.

*Forced swim test*

Rats were placed individually in Pyrex cylinders (Fischer Scientific: 21 cm × 46 cm)[2]. Cylinders contained water at 24 °C (±1 °C) up to a 30 cm mark on the cylinder. The test was a two-day experiment. In the first day, the rats were placed to the cylinder for 15 minutes for habituation. After 24 hours, the rats were placed in the same cylinders and video-taped from above for 5 minutes.

The videos were analyzed with the time-sampling technique to score the predominant behavior (immobility, climbing, or swimming) in 5-second periods by an observer blinded to the experimental groups with BORIS. Behavior was scored as immobile when the rat showed no or minimal movement, swimming with horizontal movement of the forepaws, and climbing with vertical movement of the forepaws as described previously[3]. Cumulative sums of each behavior were used in the subsequent analyses.

**RNA extraction and sequencing**

Total RNA was extracted from tissue samples using Qiagen RNeasy Mini kit following manufacturer's instructions (Qiagen, Hilden, Germany). RNA samples were quantified using a Qubit 4.0 Fluorometer (Life Technologies, Carlsbad, CA, USA) and RNA integrity was checked with an RNA Kit on Agilent 5300 Fragment Analyzer (Agilent Technologies, Palo Alto, CA, USA).

RNA sequencing libraries were prepared using the NEBNext Ultra II RNA Library Prep Kit for Illumina following manufacturer's instructions (NEB, Ipswich, MA, USA). Briefly, mRNA was first enriched with Oligo(dT) beads. Enriched mRNAs were fragmented according to manufacturer's instructions. First strand and second strand cDNAs were subsequently synthesized. cDNA fragments were end repaired and adenylated at 3'ends, and universal adapters were ligated to cDNA fragments, followed by index addition and library enrichment by limited-cycle PCR. Sequencing libraries were validated using NGS Kit on the Agilent 5300 Fragment Analyzer (Agilent Technologies, Palo Alto, CA, USA), and quantified by using Qubit 4.0 Fluorometer (Invitrogen, Carlsbad, CA).

The sequencing libraries were multiplexed and loaded on the flow cell on the Illumina NovaSeq 6000 instrument according to manufacturer's instructions. The samples were sequenced using a 2x150 Pair-End (PE) configuration v1.5. Image analysis and base calling were conducted by the NovaSeq Control Software v1.7 on the NovaSeq instrument. Raw sequence data (.bcl files) generated from Illumina NovaSeq was converted into fastq files and de-multiplexed using Illumina bcl2fastq program version 2.20. One mismatch was allowed for index sequence identification.

**Rat fecal pellet microbiota DNA extraction**

Due to availability of DNA extraction kits, different methodology was applied for the human samples. Fecal pellets were dissolved in lysis buffer and microbial DNA was extracted using a DNA Stool 200 Kit special H96 (PerkinElmer, Turku, Finland) kit with a corresponding Chemagic with Magnetic Separation Module I (MSM I) extraction robot according to manufacturer's instructions, except additional homogenization with a PowerBead Pro Plates (Glass beads 0.1 mm) and TissueLyser II (Qiagen, USA). The plate was shaken in TissueLyser II at 15 Hz for 5 minutes x2. Next, the plate was centrifuged at 4500 g for 6 min and 800 µL of lysate was transferred to the sample plate and the extraction proceeded according to the manufacturer's protocol. DNA was stored at -80°C until library preparation. The concentrations of the DNA were measured with a Qubit Fluorometer using a Qubit dsDNA High Sensitivity Assay kit (Thermo Fisher Scientific, Waltham, MA, USA). The variable region V4 of 16S ribosomal gene was amplified with custom-design dual-indexed primers[4,5]. Samples were sequenced with Illumina MiSeq (Illumina, Inc., San Diego, CA, USA) platform. The 4 nM library pool was denatured, diluted to a concentration of 4pM, and an 8% denaturalised PhiX control (Illumina, USA) was added. The library samples were sequenced with a MiSeq Reagent kit v3, 600 cycles (Illumina, USA) on a Miseq system with 2x 250 base pair (bp) paired ends following the manufacturer's instructions. A positive control, Zymobiomics Microbial community DNA standard, (Zymo Research, USA) and a negative control (PCR-grade water) were included in library preparation to control the PCR. Lysis buffer was used as negative control in DNA extraction to control contamination.

The rat 16s rRNA sequencing data was processed using R (v. 3.6.1, Vienna, Austria) and amplicon sequencing variants (ASVs) were inferred with DADA2 [24]. The reads were truncated to length 225. Reads with more than two expected errors were excluded. SILVA taxonomy database (version 138) [25] and RDP Naïve Bayesian Classifier [26] was used to assign taxonomy. DECIPHER was used for multiple sequence alignment [27].

**DNA extraction of human fecal samples**

Samples were divided into aliquots and frozen at -80°C until DNA extraction. DNA was extracted with the GXT Stool Extraction Kit VER 2.0 (Hain Lifescience GmbH, Nehren, Germany) according to manufacturer's instructions, except vortexing was replaced by homogenization with MOBIO PowerLyzer 24 Bench Top Bead-Based Homogenizer in 0.1mm glass bead tubes (MO BIO Laboratories, Inc., Carlsbad, CA, USA) at 1000 rpm for three minutes[4]. DNA concentrations were measured with Qubit dsDNA HS Assay kit and Qubit 2.0 fluorometer (Thermo Fisher Scientific, Waltham, MA, USA). DNA samples were stored at -80°C.

**Metabolome assays**

The metabolomics assays were performed as previously described [6].**Targeted BAs Measurement:** BAs were extracted by adding 40 µL of fecal homogenate to 400 µL of crash solvent (methanol containing 62.5 ppb each of the internal standards LCA-d4, TCA-d4, GUDCA-d4, GCA-d4, CA-d4, UDCA-d4, GCDCA-d4, CDCA-d4, DCA-d4, and GLCA-d4), and filtering them using a Supelco protein precipitation filter plate. The samples were dried under a gentle flow of nitrogen and resuspended using 20 µL of resuspension solution (methanol:water, 40:60). For quality control (QC) purposes, pooled QC samples and blank samples were used. Calibration curves were prepared by pipetting 40 µL of standard dilution into vials, adding 400 µL of crash solution, and drying and resuspending them in the same way as the other samples. The concentrations of the standard dilutions ranged between 0.0025 and 600 ppb. The LC separation was performed on a Sciex Exion AD 30 (AB Sciex Inc., Framingham, MA) LC system consisting of a binary pump, an autosampler set to 15 °C, and a column oven set to 35 °C. A Waters Acquity UPLC HSS T3 (1.8 µm, 2.1 × 100 mm) column with a precolumn of the same material was used. Data processing was performed using Sciex MultiQuant.

**Quantification of SCFAs:** Fecal samples were homogenized by adding water (10 µL per mg of dry weight as determined for the BA analysis) to wet feces, and the samples were homogenized using a bead beater. Analysis of SCFAs was performed on fecal homogenate (50 µL) crashed with 500 µL of methanol containing internal standards (propionic acid-d6 and hexanoic acid-d3 at 10 ppm). Samples were vortexed for 1 minute, followed by filtration using a 96-Well protein precipitation filter plate (Sigma-Aldrich, 55263-U). Retention index (RI, 8 ppm C10-C30 alkanes and 4 ppm 4,4-Dibromooctafluorobiphenyl in hexane) was added to the samples. Gas chromatography (GC) separation was performed on an Agilent 5890B GC system equipped with a Phenomenex Zebron ZB-WAXplus (30 m × 250 µm × 0.25 µm) column, and a short blank pre-column (2 m) of the same dimensions was also added. Dilution series of SCFA standards including acetic, propionic, butyric, valeric, hexanoic acid, isobutyric, and iso-valeric acid were prepared in concentrations of 0.1, 0.5, 1, 2, 5, 10, 20, 40, and 100 ppm for the construction of standard curves for quantification.

**Polar Metabolites:** Polar metabolites were extracted in methanol. The method was adapted from the method previously described Lamichhane et al. [7]. Fecal homogenate (60 µL) was diluted with 600 µL of methanol crash solvent containing internal standard (heptadecanoic acid (5 ppm), valine-d8 (1 ppm), and glutamic acid-d5 (1 ppm)). Derivatization was carried out on a Gerstel MPS MultiPurpose Sampler. GC separation was carried out on an Agilent 7890B GC system equipped with an Agilent DB-5MS (20 m × 0.18 mm (0.18 µm)) column. Mass spectrometry was carried out on a LECO Pegasus BT system (LECO). The samples were run in 9 batches, each consisting of 100 samples and a calibration curve. To monitor the run, a blank, a QC, and a standard sample with a known concentration were run between every 10 samples. Data processing was carried out using MSDIAL (version 4.7). Identification was conducted using retention index with the assistance of the GCMS DB-Public-kovatsRI-VS3 library provided on the MSDIAL webpage. Data were normalized using heptadecanoic acid as the internal standard, and the features with a coefficient of variance of less than 30% in QC samples were selected. Further filtering was performed to remove alkanes and duplicate features. The IDs of the features that passed the CV check were further verified using the Golm Metabolome Database.

### Supplemental figures

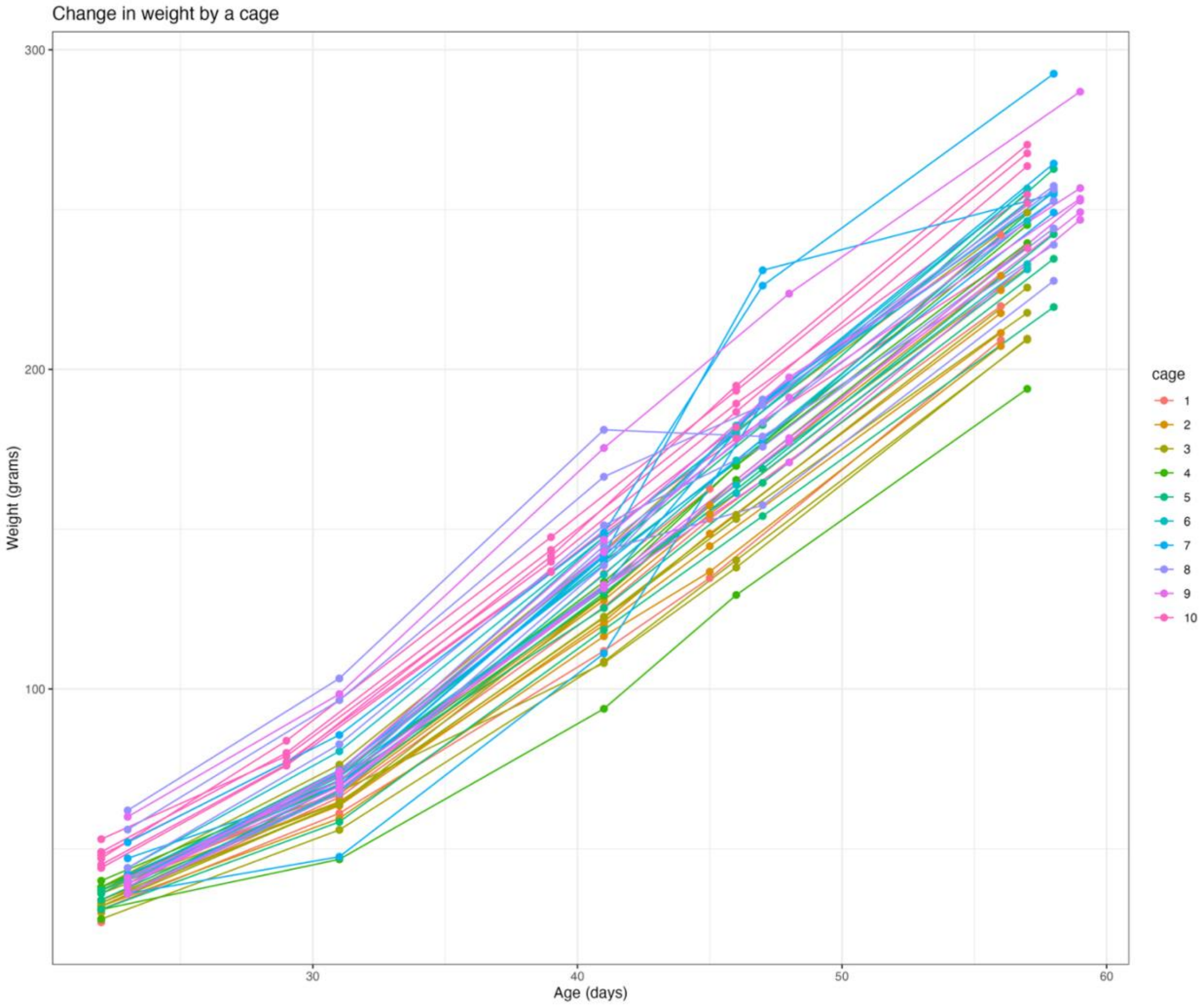


Figure S1. The change in the weight of individual rats was overall linear. Linear mixed-effect model suggested that there was an age*cage interaction (p<0.0001) in the model: *weight ~ age + cage + age * cage*.

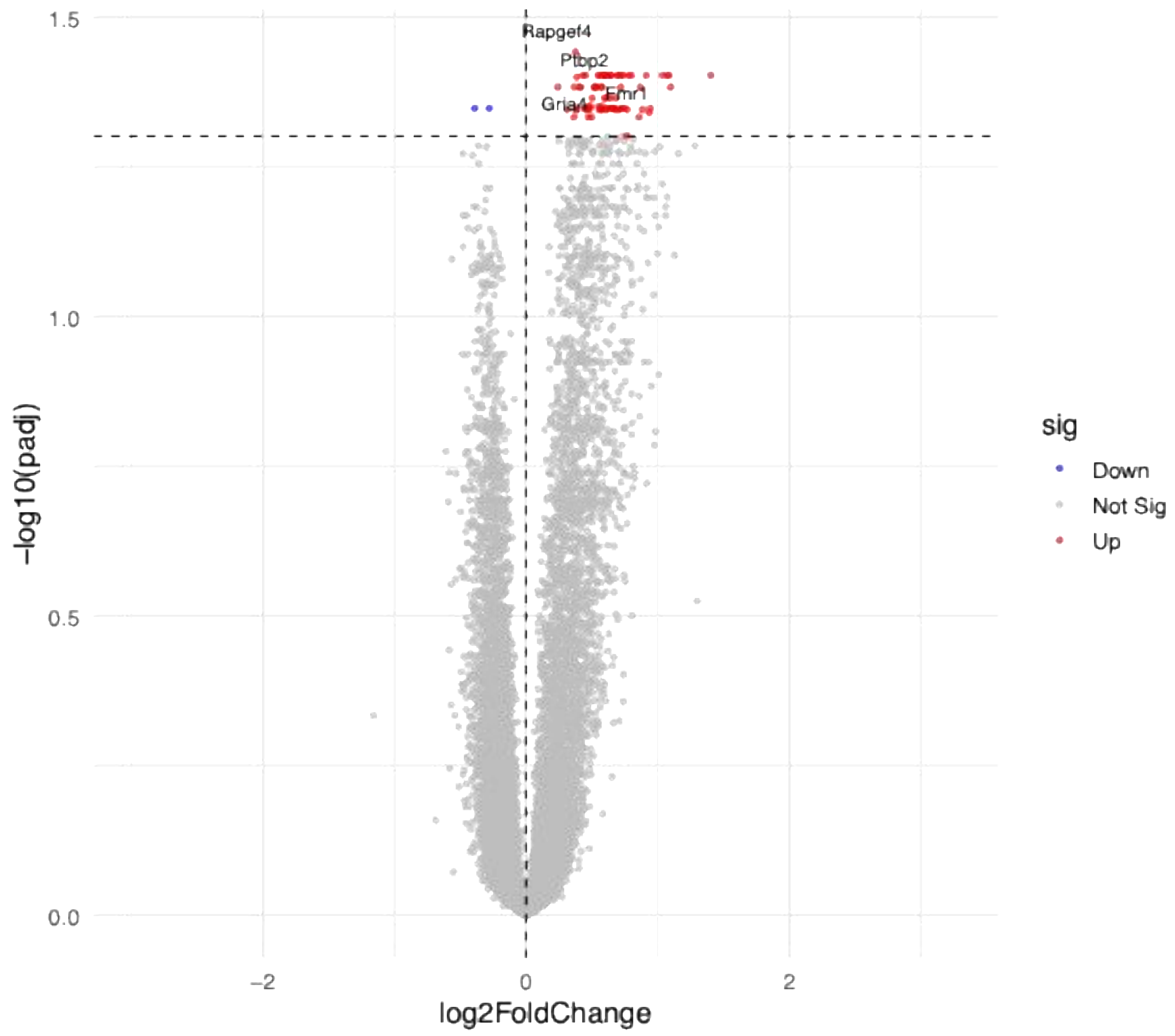


Figure S2. Volcano plot of differentially expressed genes in striata of recipients of inhibited toddlers' feces vs controls annotated with gene names related to the top significant associations.
